# Regional Microglial Response in Enthorino-hippocampal Slice Cultures to Schaffer Collateral Lesion and Metalloproteinases Modulation

**DOI:** 10.1101/2024.01.10.575060

**Authors:** A. Virtuoso, C. Galanis, M. Lenz, M. Papa, A Vlachos

## Abstract

Microglia and astrocytes are essential in sustaining physiological networks on the central nervous system, with their ability to remodel the extracellular matrix, being pivotal for synapse plasticity. Recent findings challenge the traditional view of homogenous glial populations in the brain, uncovering morphological, functional and molecular heterogeneity among glial cells. This diversity has significant implications for both physiological and pathological brain states. In the present study, we mechanically induced a Schaffer collateral lesion (SCL) in mouse enthorino-hippocampal slice cultures to investigate glial behavior, i.e., microglia and astrocytes, under metalloproteinases (MMPs) modulation in the lesioned area, CA3, and the denervated region, CA1. We observed distinct response patterns in microglia and astrocytes 3 days after the lesion. Notably, GFAP-expressing astrocytes showed no immediate changes post-SCL. Microglia responses varied depending on their anatomical location. The MMPs inhibitor GM6001 did not affect microglial reactions in CA3, while increasing the Iba1 cells numbers in CA1, underscoring the complexity of the hippocampal neuroglial network post-injury. These findings highlight the importance of understanding glial regionalization following neural injury and MMPs modulation, and pave the way for further research into glia-targeted therapeutic strategies for neurodegenerative disorders.

## 1. Introduction

Microglia, as integral components of the resident innate immune defense in the central nervous system (CNS), work alongside astrocytes in maintaining physiological networks. Their pivotal role is increasingly recognized in various neurological disorders. Recent studies challenges the traditional perspective of an homogenous glia population in the brain, revealing both morphological and molecular diversity among glial subpopulations, even in resting states [1]. The heterogeneity of glial morphology is mirrored by diverse functional capabilities, adding complexity to our understanding. This necessitates further investigation, particularly for the advancement of drug discovery and the development of personalized medicine approaches.

The phenotyping of astrocytes and microglia in CNS disease reveals distinct patterns based on their anatomical locations, as demonstrated through transcriptional, genetic, morphological, and metabolic activities [2] [3] [4]. Furthermore, the sensome of human microglia from various brain regions exhibits unique reactions to inflammatory stimuli, which change rapidly [5]. This introduces time as a critical variable in such studies, underscoring the dynamic nature of glial responses in the context of CNS disorders.

The activity of microglia and astrocytes, encompassing proliferation, motility, phagocytosis and synaptic pruning, is influenced by the remodeling of the extracellular matrix (ECM), which is critical beyond the factors of origin, trigger, and timing of pathological processes [6]. Metalloproteinases (MMPs), zinc-dependent enzymes, are mainly responsible for the structural reorganization of the CNS during development, repair, and plasticity. The levels of MMPs are regulated by the components of the neurovascular unit, balancing between adaptive and maladaptive plasticity [7]. This regulation plays a significant role in determining the progression and severity of neurological disorders [8].

Modulation of MMPs has shown promising results in improving deficits in various disease models. In Alzheimer’s disease models MMPs modulation has led to notable improvements [9]. Similar beneficial effects have been observed in peripheral nerve injury [10,11] and reducing tumor metastasis [12]. However, the impact of MMPs on hippocampal synaptic plasticity presents inconsistent findings [13,14]. To understand the cellular and molecular mechanisms involving glial cells plasticity, various functional and mechanical lesion models have been established [15–18]. Despite these advancements, the precise links and underlying mechanisms connecting reactive gliosis, MMPs modulation, and the progression of neurological disorder remain to be fully elucidated.

In this study, we mechanically induced a Schaffer collateral lesion (SCL) in mouse enthorino-hippocampal slice cultures. This approach was used to characterize microglia and astrocytes under MMPs modulation in the lesioned area, CA3, and the denervated region, CA1. We analyzed adaptations of both microglia and astrocytes. Intriguingly, our findings revealed heterogenous responses of glial cells in CA3 and CA1. These results reveal the significant influence of anatomical location, the nature of the stimulus, and the timing of the disease process on glial cell behavior, and hence the complex dynamics of glial cell responses in CNS pathologies.

## 2. Materials and Methods

### Ethics statement

Mice were maintained in a 12h light/dark cycle with food and water available ad libitum. All experimental procedures were performed according to the German animal welfare legislation and approved by the animal welfare committee and/or the animal welfare officer at the University of Freiburg, Faculty of Medicine (X-17/07K, X-18/02C, X-21/01B). All the procedures were conducted minimizing distress and pain to the animals.

### Preparation of enthorino-hippocampal organotypic tissue cultures

Entorhino-hippocampal tissue cultures were prepared at postnatal day 4–5 from C57BL/6J mice of either sex as previously described [19]. Cultivation medium contained 50% (v/v) MEM, 25% (v/v) basal medium eagle, 25% (v/v) heat-inactivated normal horse serum, 25mM HEPES buffer solution, 0.15% (w/v) bicarbonate, 0.65% (w/v) glucose, 0.1mgml^−1^ streptomycin, 100Uml^−1^ penicillin, and 2mM glutamax. The pH was adjusted to 7.3 and the medium was replaced three times per week. All tissue cultures were allowed to mature for at least 18 days to ensure the stability of structural and functional properties of the organotypic tissue [20–23]. Entorhino-hippocampal tissue cultures were maintained in a humidified atmosphere with 5% CO2 at 35°C.

### Schaffer Collateral lesion (SCL)

Mechanical pathway transection was performed with a sterile scalpel in mature tissue cultures (≥ 18 days in vitro) as described in [18]. SCL was applied between the CA3 and the CA1 region of the hippocampus, without affecting the perforant path projections to CA3 (Fig.1a). Except for the lesion-induced partial denervation of CA1 pyramidal neurons, cytoarchitecture of both the hippocampus and the entorhinal cortex remained unchanged (Fig.1a).

**Figure 1.**
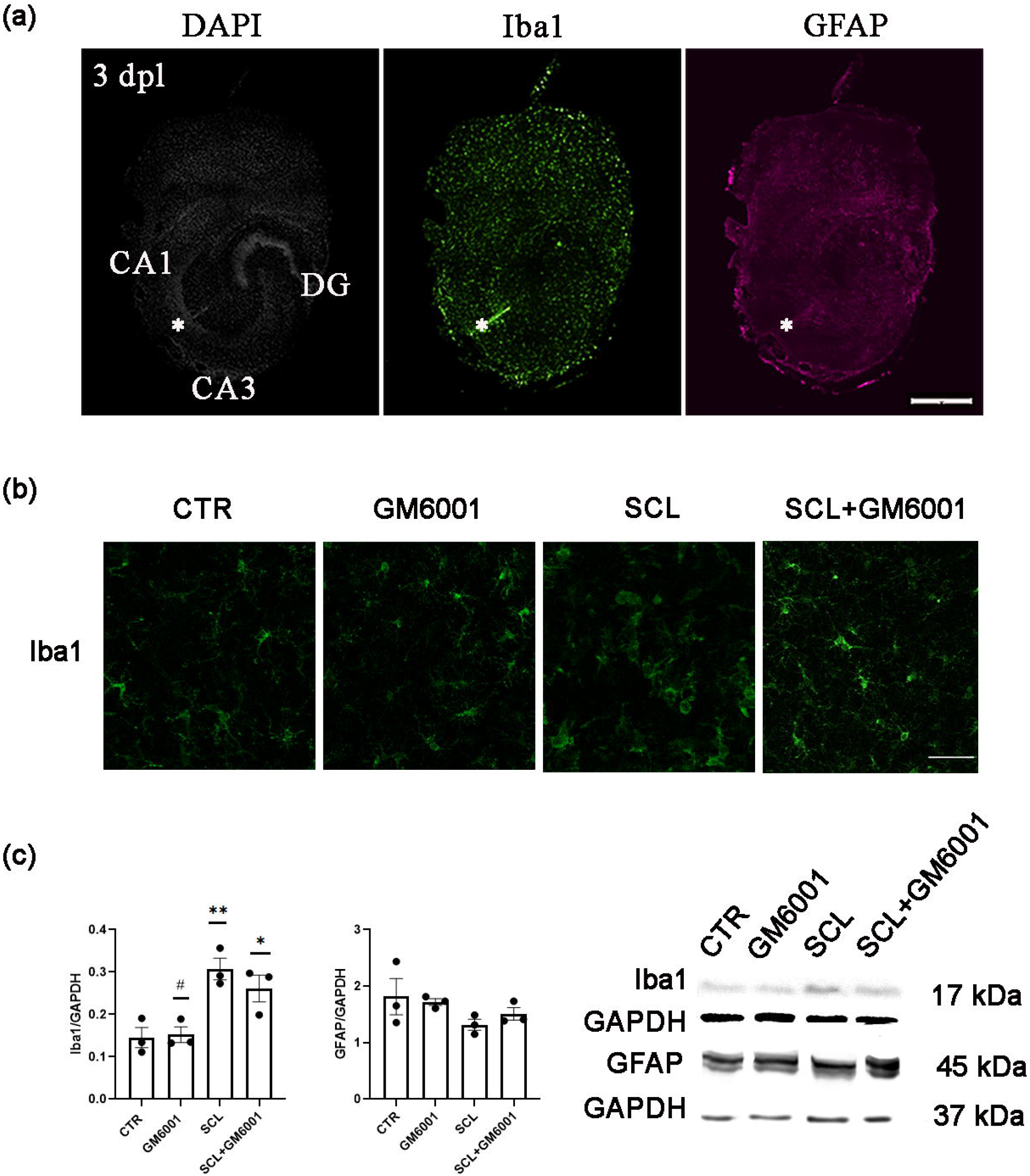
Microglia is activated three days after Schaffer collateral lesion (SCL) in vitro. (a) Representative enthorino-hippocampal slice culture stained for DAPI (white), Iba1 (green) and GFAP (magenta) 3 days after lesion (3dpl). The SCL between hippocampal areas cornu ammonis 3 (CA3) and cornu ammonis 1 (CA1) is indicated with asterisks. Scale bar 400µm. (b) Representative image of Iba1 positive cells at the lesion site from CTR, GM6001, SCL and SCL+GM6001 groups. Scale bar 50 µm. (c) Western blot relative quantification and corresponding bands for Iba1 (left) and GFAP (right) from whole control (CTR) and lesioned (SCL; 3dpl) enthorino-hippocampal slice cultures. A set of slice cultures were treated with the broad-spectrum MMP inhibitor GM6001 (50 nM; n = 3 cultures per group).

### Pharmaceutical treatment for MMPs modulation

MMPs modulation was performed using the broad-spectrum MMPs inhibitor, GM6001, Ilomastat (Merck, Millipore). The GM6001 was prepared as per manufacturer instructions and dissolved in DMSO. GM6001 was then added in the slice cultures medium at a final concentration of 50 nM as a pretreatment, 1 h before the SCL. Fresh GM6001 was added every time the medium was replaced.

### Experimental groups

Slices cultures were divided into four groups: (I) Control, receiving 1 uL of DMSO in the fresh cultivation medium; (II) GM6001, having 1 uL of GM6001 50 uM in the fresh cultivation medium; (III) SCL, subjected to the Schaffer collateral lesion (SCL); (IV) SCL+GM6001, receiving the lesion after 1 hour of pre-treatment with GM6001.

### Immunofluorescence (IF)

At 3 days post-lesion (dpl), the slices were processed for immunostaining. Cultures were fixed in a solution of 4% (w/v) paraformaldehyde (PFA) in phosphate-buffered saline (PBS, 0.1M, pH 7.4) and 4% (w/v) sucrose for 1h. Fixed cultures were incubated for 1h with 10% (v/v) normal goat serum (NGS) in 0.5% (v/v) Triton X-100-containing PBS to block non-specific staining. Whole tissue cultures were incubated with rabbit anti-Iba1 (1:1000; Fujifilm Wako, #019–19741) or mouse anti-GFAP (1:1000; Sigma) in PBS containing 10% (v/v) normal goat serum (NGS) and 0.1% (v/v) Triton X-100 at 4°C overnight. Cultures were washed and incubated for 3h with appropriate secondary antibodies (Alexa Fluor anti-rabbit 488; Alexa Fluor anti-mouse 555, 1:1000, in PBS with 10% NGS or NHS, 0.1% Triton X-100; Invitrogen). All the nuclei were visualized by DAPI staining (1:5000 in PBS for 10min; Thermo Scientific, #62248). Sections were washed, transferred onto glass slides and mounted for visualization with anti-fading mounting medium (DAKO Fluoromount).

Confocal images in immunostainings were acquired using a Leica SP8 confocal microscope equipped with a 20x (NA 0.75, Leica) or 40x (NA 1.3, Leica) objective lens. Detector gain and amplifier were initially set to obtain pixel intensities within a linear range.

### Western blotting (WB)

At 3 dpl, the slices were processed for western blotting. Whole enthorino-hippocampal slices cultures were homogenized in RIPA buffer enriched with Phenylmethanesulfonyl Fluoride (PMSF) and protease inhibitors cocktail. The protein content of each sample was determined using the Bradford method and the same amount of proteins was loaded in polyacrylamide gels (10-15 %). The proteins were separated during the SDS page and transferred to a PVDF membrane using the Biorad Tetra System. After blocking in 5% non fat milk in T-TBS (Tris buffer phosphate+Tween) for 35 min, the membranes were incubated with the primary antibody against GFAP (mouse, 1:1000, Sigma), Iba1 (rabbit, 1:500, Wako) and GAPDH (rabbit, 1:500, Novus-Bio) at 4°C. After incubation with the proper secondary antibody, detection of the targets was performed using chemiluminescence. The expression of Iba1 and GFAP band was normalised to the expression of the corresponding loading control, GAPDH. WB experiments were repeated at least three times.

### Quantification and statistics

All the quantifications were performed using Image J software. Semi-automated procedure was conducted to quantify non laminar microglia hypertrophy and cell count in CA1 and CA3 regions. The hypertrophy index was assessed as percentage (%) of area covered by the Iba1+ cells compared to the total scanned area in CA1 and CA3 using a fixed ROI (region of interest) spanning from the stratum oriens to the stratum moleculare. The Iba1+ cell count was expressed as % relative to the number of cells identified in CA3 and CA1 of the CTR group, respectively. Immunostainings files were re-named using a code and analysis were conducted blinded.

Data were statistically analyzed using GraphPad Prism 9 (GraphPad software, USA). Normal (Gaussian) distribution was tested using Shapiro-Wilk test. For statistical comparison of data sets with three or more experimental groups passing the normality test, one-way ANOVA test with Tukey’s correction for multiple comparisons was used. For the evaluation of data sets with three or more experimental groups not disributed under Gaussian curve, a Kruskal-Wallis test followed by Dunn’s posthoc correction was applied. p-values <0.05 were considered as statistically significant (*p<0.05, **p<0.01, ***p<0.001). The n-numbers are provided in the figure legends. (*) was used to show the significant differences relative to the CTR group, while (#) was used when detected through multiple comparisons among the experimental groups. In the text and figures, data are expressed as mean±standard error of the mean (s.e.m.).

### Digital illustrations

Confocal image stacks were stored as TIF-files. Figures were prepared using the ImageJ software package3 and Adobe Photoshop graphics software. Images were adjusted for brightness, contrast and sharpness, mantaining the same parameters among the groups.

## 3. Results

### 3.1 Iba1+ cells respond to the SCL at 3 dpl, unlike GFAP-expressing astrocytes

SCL was performed in mature (18 div) enthorino-hippocampal tissue cultures prepared from 4-5 days old mice, using a sterile scalpel. Three days after SCL, a notable expression of Iba1 positive cells, indicative for microglia, was observed around the lesion site. Conversely, GFAP immunostaining, a marker for astrocyte activation, remained unchanged at the lesion site (Figure 1a), implying a differential glial response to the injury.

Further examination using high magnification lens confirmed the increase in Iba1-responsive elements at the lesion site. Interestingly, treatment with GM6001 ameliorated the microglial reaction (Figure 1b).

Western blot analysis conducted on the whole slices revealed a significant increase in Iba1 expression in the SCL group, while GFAP levels did not show any significant differences among the experimental groups (Figure 1c). Notably, Iba1 overexpression was modestly reduced following GM6001 [50 nM] treatment, although this decrease was not statistically significant (Figure 1c).

These data suggested that SCL selectively triggers a microglia response at early stages post-injury, without significantly impacting reactivity of GFAP-expressing astrocytes at 3 dpl.

### 3.2 Iba1+ cells show a region-specific response to the SCL in enthorino-hippocampal slices cultures

Iba1 protein overexpression is a key feature of reactive gliosis [24], potentially resulting from cell hypertrophy and/or increased microglia density. To investigate this, hypertrophy measurements and cell counts were conducted in a representative area of CA3, located 0,5 mm from the lesion site, and in a symmetrical region in CA1, as depicted in Figure 2a. Iba1 immunostaining effectively highlights both cell body and processes, facilitating the assessment of morphological diversity among myeloid cells [25]. This makes Iba1 a valuable marker for quantifying the extent of microglial activation, allowing for both the measurement of the area occupied by Iba1+ cells, serving as a hypertrophy index, and cell counting as shown in Figure 2b and 2c.

**Figure 2.**
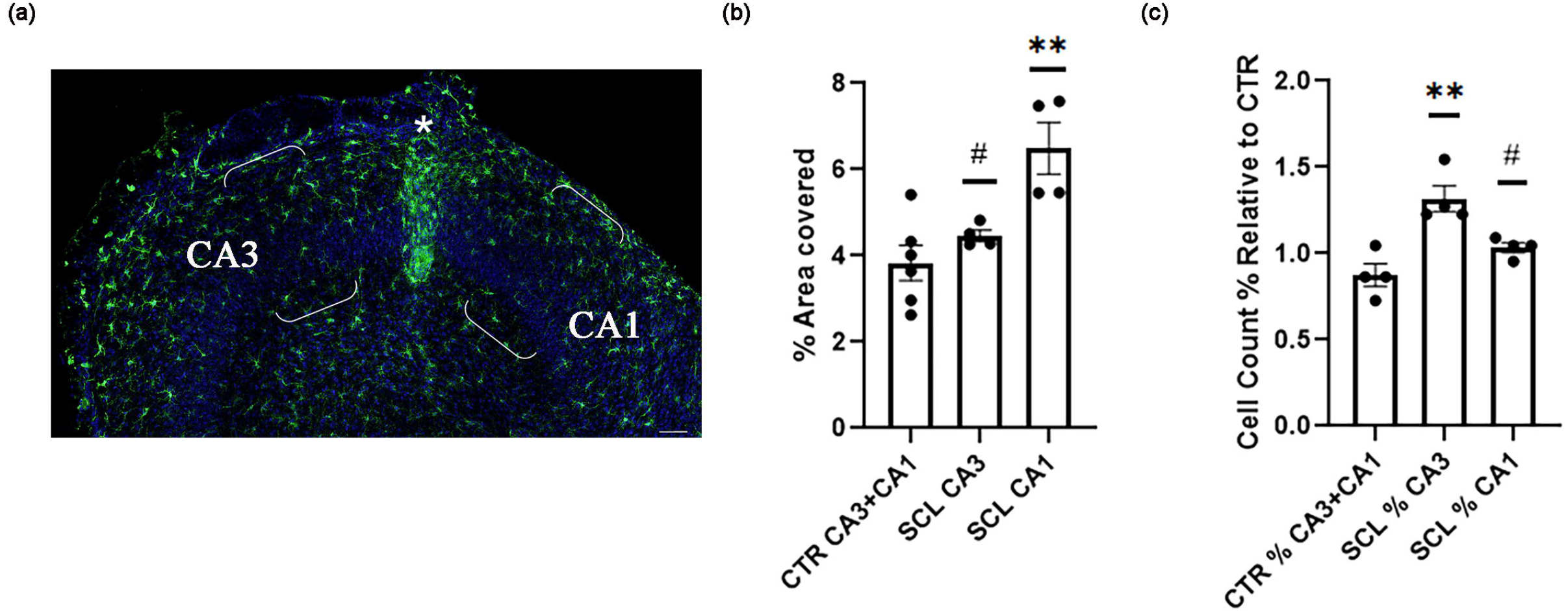
Region-specific effects of SCL on microglia. (a) Hippocampal slice culture stained for Iba1 (green) and DAPI (blue). Microglia was analyzed in specified regions of CA3 and CA1, 0.5 mm lateral to the lesion site (marked with an asterisk). Scale bar 200 µm. (b) Data analysis of % area covered by Iba1 expressing cells relative to the total scanned area (n = 4-6 cultures per group). (c) Iba1 positive cells count relative to the cell count in the pooled CTR group (%) (n = 4 cultures per group).

The analysis revealed a notable difference in the response of Iba1+ expressing cells across regions. Specifically, the area covered by Iba1+ cells in CA1 was significantly larger compared to both the control (CTR) group and CA3 (Figure 2b). Interestingly, this increase in coverage did not correspond to a significant increase in cell count in CA1. In contrast, microglia in CA3 did not exhibit hypertrophy when compared to the CTR group, yet there was an observable increase in the their number (Figure 2c).

Taken together, these data highlight a distinct regionalization in the response of Iba1-expressing cells to the SCL. The regional variation points to a nuanced heterogeneity, possibly attributed to the differential nature of the stimulus: direct nerve damage in CA3 and denervation to CA1. This suggests that the microglial response is finely tuned not only to the presence of injury but also to the specific type of neural insult.

### 3.3 MMPs modulation lead to a re-organization of the CA1 microglia response to SCL

Microglia activities such as proliferation, hypertrophy, and motility within the neuro-glial network require precise remodeling of the ECM, largely driven by MMPs activity [8]. Notably, MMPs are implicated in hippocampal synaptic plasticity in both CA3 and CA1 regions, although the underlying molecular mechanisms may differ significantly [26]. Furthermore, CA1 and CA3 exhibit notable differences in their transcriptional and proteomic profiles under basal conditions [27,28].

In this context, treatment with the MMPs-broad spectrum inhibitor GM6001 [50 nM] for 3 days did not affect microglia in CA3, both under control conditions and 3 dpl (Figure 3a-b). In contrast, with GM6001 treatment, CA1 microglia maintained hypertrophic characteristics associated with SCL and exhibited an increased number compared to the CTR group (Figure 3a-c).

**Figure 3.**
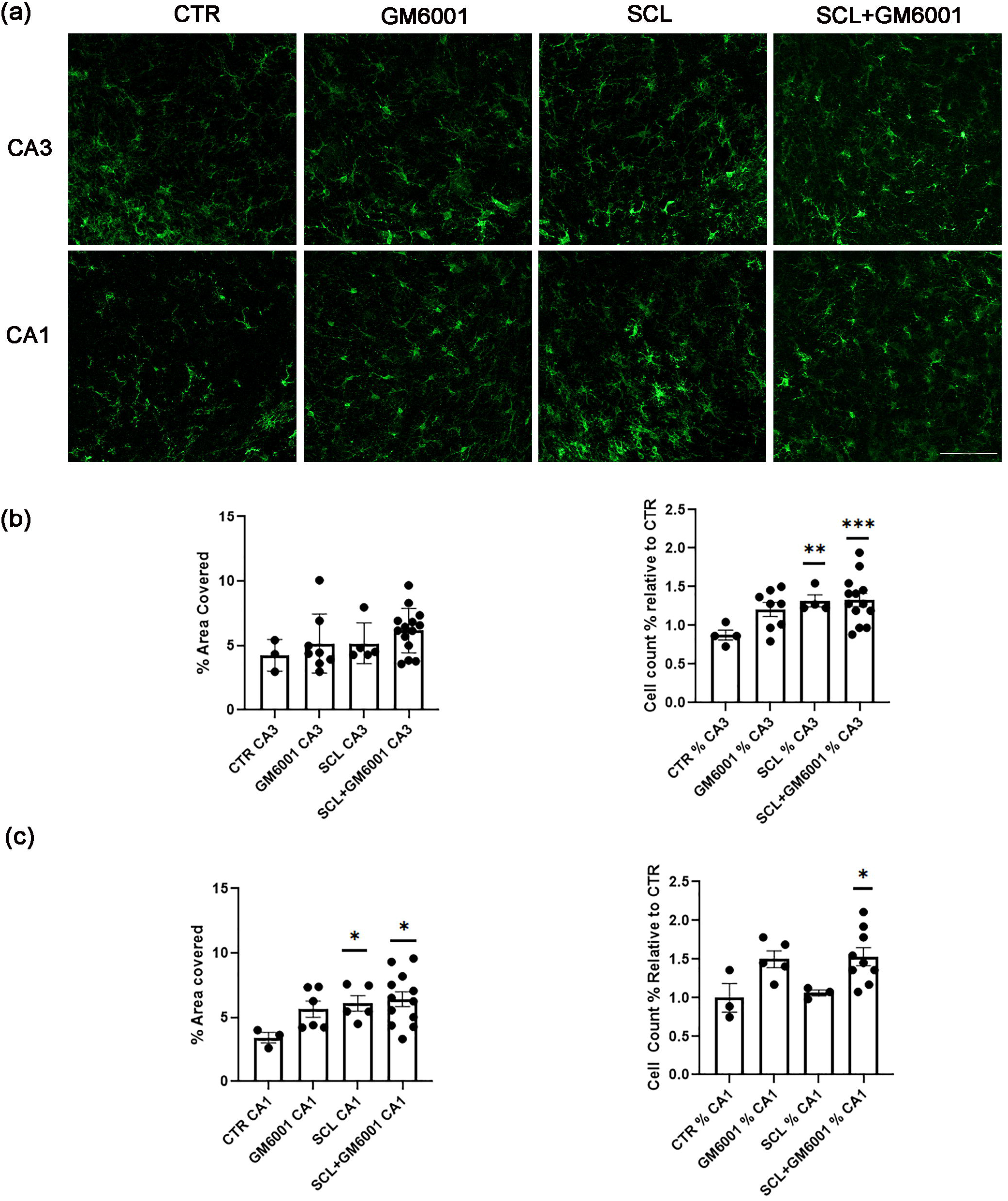
Effects of metalloproteinases on SCL-induced microglia responses. (a) Representative images of Iba1 positive cells from CTR, GM6001, SCL, and SCL+GM6001 groups. Scale bar 100µm. (b, c) Analysis of % area covered by Iba1 expressing cells relative to the total scanned area (left), and cell count expressed in % relative to the CTR (right) (CA3, n = 4-13 cultures per group; CA1, n = 3-12 cultures per group).

These results support the concept of region-specific microglial responses following MMPs modulation, emphasizing the complexity and diversity of glial plasticity. They highlight the importance of a detailed understanding of regional and molecular variations in glial cell responses within the CNS.

### 3.4. Figures

## 4. Discussion

CNS plasticity has been a subject of extensive study, with the hippocampus often serving as a primary model due to its distinct structure. Encased by archicortical and neocortical structures, the hippocampus can mimic a “closed” network of higher order providing a unique environment for studying neural dynamics and plasticity. Enthorino-hippocampal slice cultures are particularly valuable in this context as they allow for in vitro investigation while preserving the complex cytoand fiberarchitecture of the neuroglial network. This makes them an effective experimental model for studying the morphological, molecular and functional dynamics of glial cells [18,19,29].

Glial cells are increasingly recognized for their pivotal role in various neurological disorders, including brain trauma, spinal cord injury, and neurodegenerative diseases [30,31]. Yet, a comprehensive understanding of glial plasticity, encompassing for the nature of the trigger, the timing of the reactive gliosis, and the specificity of the brain region involved, remains an area in need of further research [32].

In this study, we utilized enthorino-hippocampal slice cultures and mechanically induced a lesion to transect Schaffer collaterals connecting CA3 to CA1. Three days after SCL, there was a marked increase in Iba1 protein expression in the hippocampal lysate, indicating an early microglial response, with no simultaneous alterations in GFAP levels, suggesting that astrocytes did not activate to the same extent initially. This suggests that microglia are swift to detect lesion-induced changes within the neuroglial network and may act as initial responders in reactive gliosis [33,34].

Our experiments further highlight the regional specificity of the microglial responses to SCL in CA3 and CA1. In CA3 microglia showed an increase in density, suggesting that cell migration and/or proliferation occur as rapid responses to direct neural injury [35]. In contrast, CA1 microglial cells exhibited hypertrophy, a feature of activated microglia characterized by enlarged somata and processes, and sometimes a reduction in ramification, distinct from ameboid, phagocytic fully activated microglia [36]. The denervation of CA1 neuron may require highly coordinated modifications in the neuroglial network compared to a direct damage of CA3 axons, especially considering that (1) CA1 dendritic spine density drops early after the lesion [37,38], (3) dendrites retract [39], (3) the transected CA3 axons degenerate and are cleared by glial cells in the denervated region [40,41].

The proliferation, migration, and activation of microglia in the CA3 region seems not to depend on MMPs-mediated ECM remodeling during the early phase after SCL, since microglial morphology and counts were unaffected by GM6001-induced modulation of MMPs. Previous research on spinal cord networks revealed elevated MMPs levels following peripheral nerve injury [42,43]. Additionally, it has been found that reactive gliosis can be reduced by inhibiting MMPs at later stages after injury [11,44]. Hence, the MMPs-dependent remodeling of ECM in CA3 may account for changes seen in later stages, such as postlesional axonal sprouting [45].

In CA1, administering GM6001 resulted in increased microglia density while preserving hypertrophy, indicating that MMPs influence microglia function in denervated brain regions during the early post-lesion phase. The biological significance of these alterations is yet to be determined. Earlier research showed denervation-induced spine density changes in CA1, but no significant functional alterations in CA1 neurons post-SCL at 3 days [18]. Transcriptomic analysis in this previous study also revealed higher MMP16 mRNA levels post-SCL [18], a membrane-bound MMP expressed by microglia, potentially involved in synaptic rewiring and hippocampal spine density regulation [18,46,47]. Regardless of these considerations, the results of the present study clearly show distinct responses in glial cells following SCL and with GM6001, a broad-spectrum MMP inhibitor.

## 5. Conclusions

The present study highlights the significance of region-specific localization in the CNS with respect to glial cell behavior. It shows that glial responses can differ based on the specific CNS region involved, the nature of the initiating event, and the timing of neural plasticity. Enhancing our understanding of the region-specific (and time-dependent) dynamics of glial cells in pathological states can provide deeper insights into both adaptive and maladaptive processes. Such knowledge is crucial for developing targeted glia-specific therapeutic strategies for neurodegenerative disorders.

## Supporting information

Representative original blots for Iba1 and corresponding internal control GAPDH.

Representative original blot for GFAP and corresponding internal control GAPDH.

## Author Contributions

Conceptualization, A.Vi., C.G., M.P. and A.V.; methodology, M.L.; formal analysis, A.Vi.; investigation, A.Vi.; resources, A.V.; data curation, A.Vi.; writing—original draft preparation, A.Vi.; writing—review and editing, C.G., M.L., A.V.; visualization, A.Vi.; supervision, M.P. and A.V.; project administration, A.V.; funding acquisition, M.P and A.V. All authors have read and agreed to the published version of the manuscript.

## Supplementary materials

Fig. S1-S2. Representative original blots for Iba1, GFAP, and their corresponding internal control GAPDH, respectively.

## Data Availability Statement

Data are available from the authors under reasonable request.

## Acknowledgments

We would like to thank Monika Paetzold for excellent technical support.

