## Supplementary figures and images for "Regional Microglial Response in Enthorino-hippocampal Slice Cultures to Schaffer Collateral Lesion and Metalloproteinases Modulation"

### Representative original blot for GFAP and corresponding internal control GAPDH.

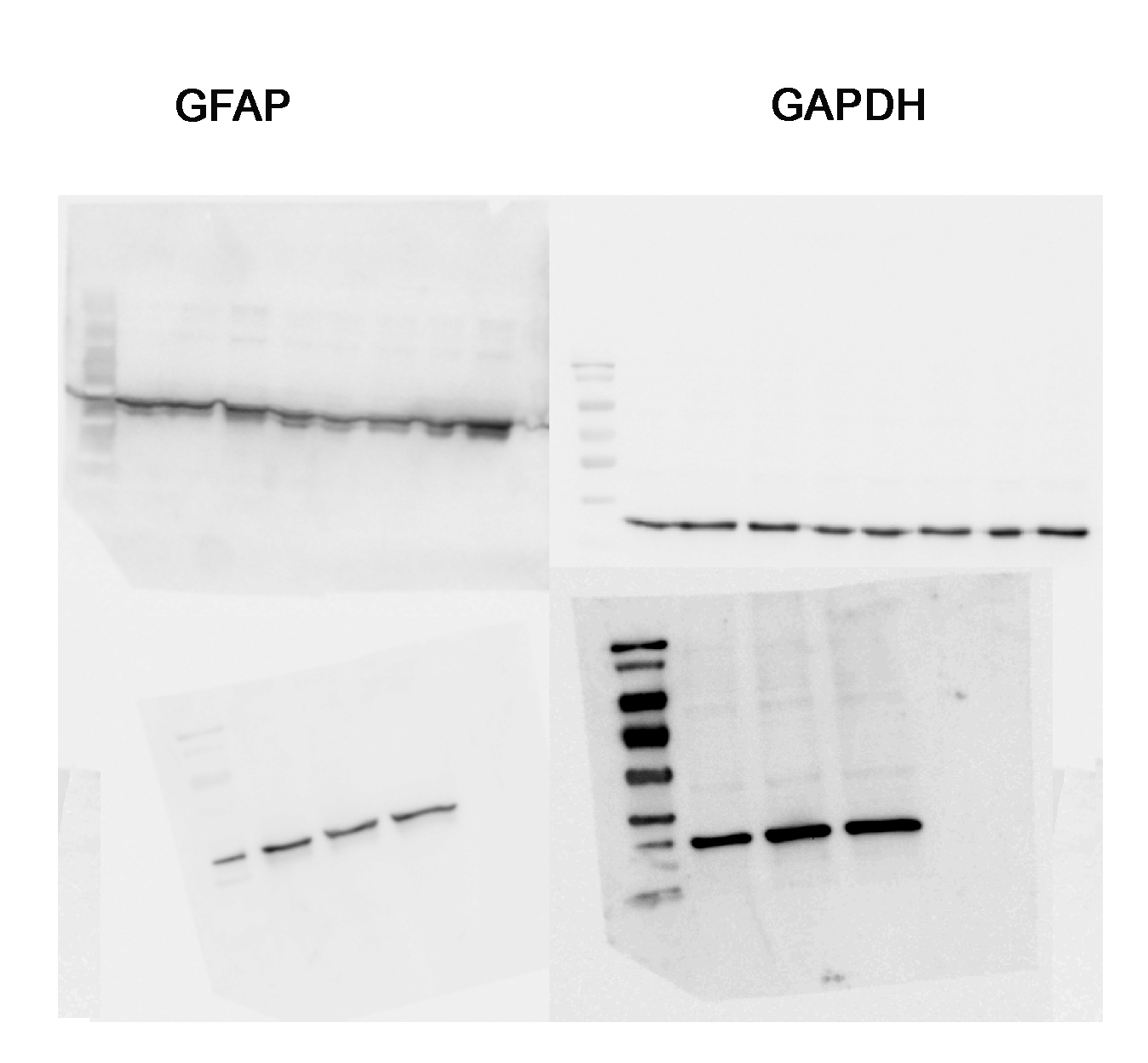

### Representative original blots for Iba1 and corresponding internal control GAPDH.

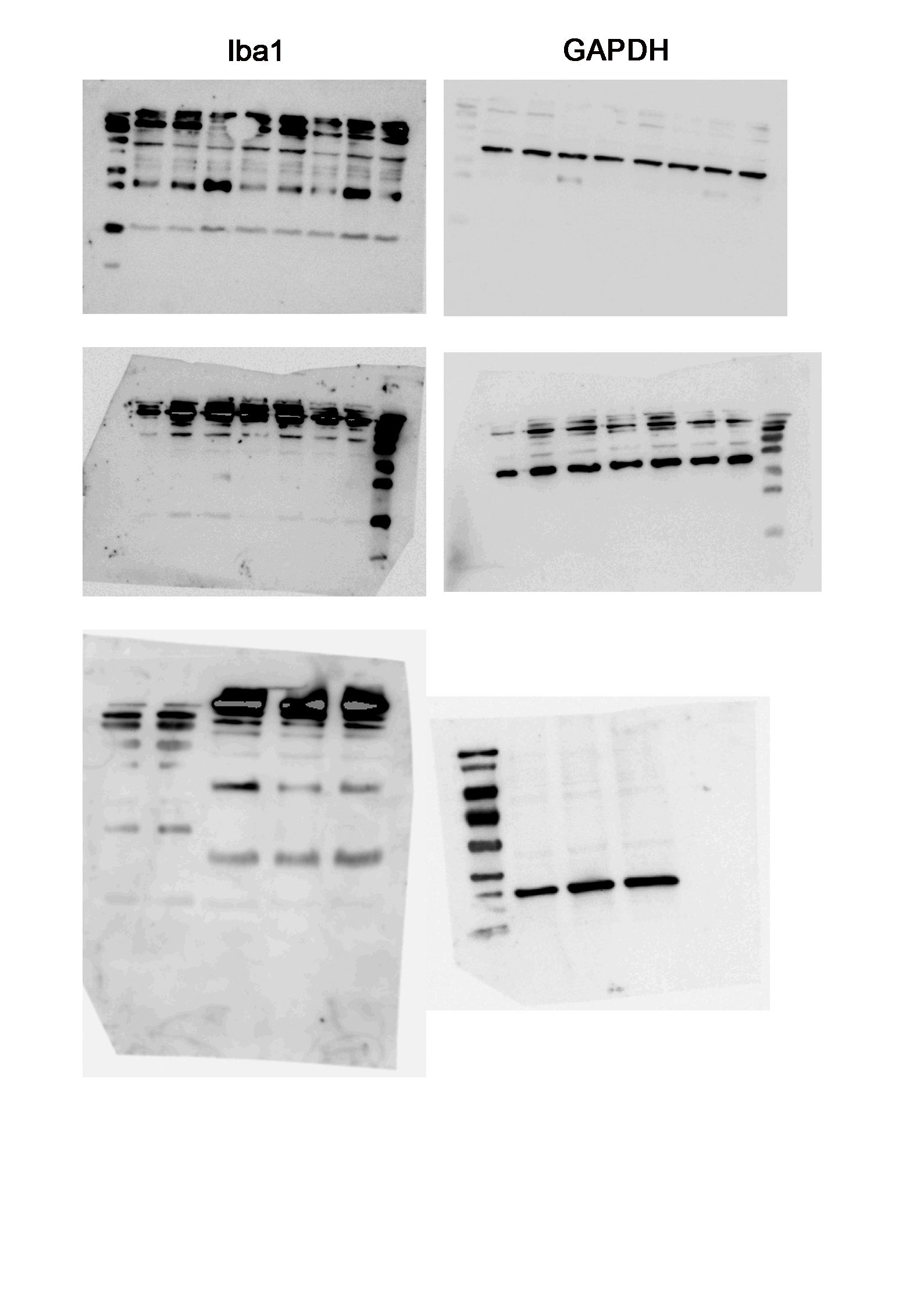
